# Persistent activity of cerebellar molecular layer interneurons facilitates anticipatory control of tongue movements

**DOI:** 10.64898/2026.01.25.701635

**Authors:** Cedric Galetzka, Mohamed Eltabbal, Bernd Kuhn

## Abstract

Precise, anticipatory movements depend on the brain’s ability to generate motor commands in advance of immediate sensory input. Although the cerebellar cortex receives abundant sensory input via the mossy-fiber pathway, the mechanisms by which continuous sensory signals are transformed or gated remain poorly understood. Here, we show that molecular layer interneurons acquire persistent sensory-related Ca^2+^ activity in mice that develop anticipatory behavior. Employing a targeted lick-interception task, we found that interneurons in lobule Crus I, but much less in Crus II, exhibit enhanced Ca^2+^ activity up to five seconds before movement onset, locked to the spatial position of a continuously moving target, independent of body movement. In more anticipatory mice, movement onset becomes linked to the dynamics of this persistent activity, whereas less anticipatory mice adjust their movements based on immediate sensory-related input. Our results indicate that anticipation in sensorimotor tasks is supported by dynamic sensory representations encoded by interneurons.

## Introduction

Anticipatory behavior requires animals to produce motor commands before immediate sensory input (Wolpert et al., 1998). The cerebellum plays an essential role in anticipatory control by building internal models that replicate the input-output relationships between the body and its environment (Kawato, 1999). These models allow the cerebellum to simulate the consequences of movement, thereby facilitating rapid feedforward motor control (Ebner, 2013; Shadmehr & Mussa-Ivaldi, 1994; Wolpert et al., 1998).

Within the cerebellar cortex, the primary output cells, Purkinje cells, refine sensorimotor associations to produce predictive motor commands by integrating two streams of information: Climbing fiber and parallel fiber input (Hull & Regehr, 2022; Medina et al., 2000; Roome & Kuhn, 20218; 2020). Climbing fibers originate from olivary neurons and provide powerful excitatory input thought to convey instructive signals related to reward, sensory salience, or error (Bina et al., 2021; Kitazawa et al., 1998; Marr, 1969; Sendhilnathan et al., 2021). Parallel fibers, the axons of granule cells that integrate multimodal sensorimotor and even cognitive input from mossy fibers, also provide excitatory input to Purkinje cells as well as molecular layer interneurons (MLIs; Huang et al., 2013; Ishikawa et al., 2015; Hull & Regehr, 2022; Wagner et al., 2017). Plasticity at parallel fiber-Purkinje cell synapses guided by climbing fiber activity is regarded as central to sensorimotor learning, such as in eyelid conditioning (Ito, 1970; Marr, 1969; Mauk & Buonomano, 2004; Kawato et al., 2021; Roome & Kuhn, 2018; 2020; Wang et al., 2000).

The role of Purkinje cells in predictive control is well-established, yet significant gaps remain in our understanding of cerebellar processing of sensory stimuli. First, although granule cells exhibit second-long ramping activity associated with reward expectation or sensory stimulation, previous studies have predominantly employed operant-conditioning task designs and static or discrete stimuli (Garcia-Garcia et al., 2024; Prat et al., 2024). However, a critical question is how continuous input in dynamic environments is represented and maintained in the cerebellar cortex. Second, while MLIs provide inhibitory input to Purkinje cells capable of modulating climbing-fiber-evoked complex spikes, their precise contribution to feedforward control of movements remains poorly understood (Gaffield & Christie, 2017; Rowan et al., 2018). Addressing these gaps is essential for understanding how the cerebellum may anticipate sensory input and provide predictive motor control.

In this study, we employed our previously developed targeted lick-interception task (Eltabbal et al., submitted), in which mice intercept moving food pellets with their tongue, combined with wide-field Ca^2+^ imaging of MLIs in Crus I and II. Inspired by the finding that MLIs encode sensory valence (Ma et al., 2020), we hypothesized that they also encode dynamic aspects of salient sensory cues that guide anticipatory tongue movements.

## Results

### Targeted lick-interception task

We studied skilled licking behavior by training head-fixed mice for up to eight consecutive days to intercept approaching food pellets by extending and retracting their tongue (Figures 1a). Trials were classified as ‘hit’ or ‘miss’ based on whether the mice grasped the pellet from the stage (Figure S1A). Trials with no detected licks were classified as ‘no lick’. The speed of the pellet on each trial varied randomly in discrete steps from 1 to 7 cm/s (step size: 1 cm/s). Early in learning, mice successfully intercepted food pellets at slower speeds (1-2 cm/s), but performance at faster speeds (3-7 cm/s) showed deviations from optimal performance (Figure 1b). With continued training, behavioral performance for faster speeds improved and the average number of licks decreased (Figure S1b). Therefore, the first and last three sessions were classified as ‘early’ and ‘late’ learning, respectively.

**Figure 1:**
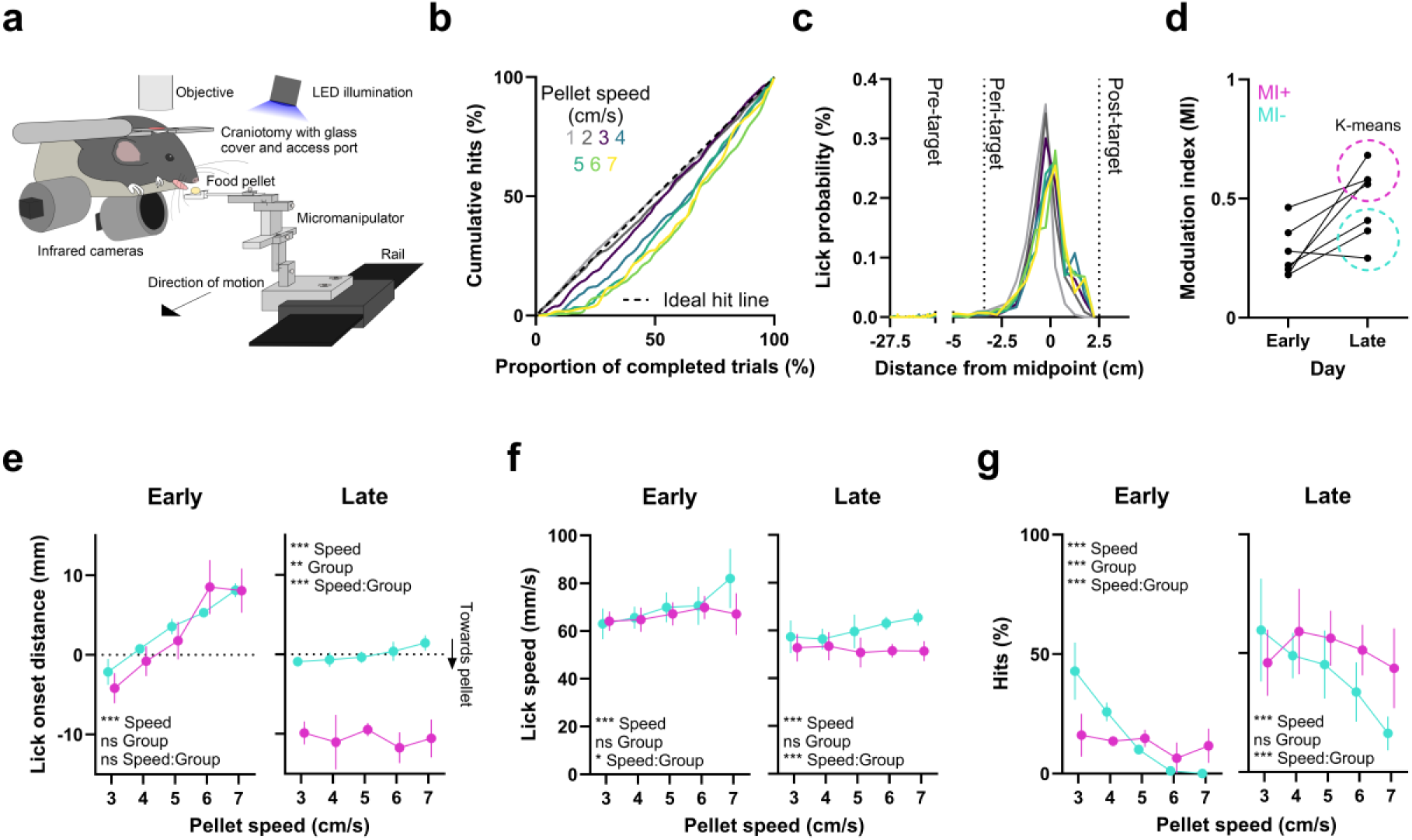
Mice differentially modulate lick onset distance and lick speed with learning. **(a)** Task setup. White noise was played to mask mechanical noise during pellet movement. **(b)** Normalized behavioral performance for all pellet speeds. Results were first computed for individual mice and then averaged across mice (N = 7 mice). **(c)** Probability distributions of lick onsets for all pellet speeds. Vertical lines separate the pre-, peri-, and post-target intervals. Data were first computed for individual mice and then averaged across mice (N = 7 mice). **(d)** Lick modulation index for all mice during early and late learning. Circles indicate K-means cluster assignments, retroactively assigned to early learning as well. **(e)** Left: Lick onset distance for pellet speeds 3-7 cm/s during early learning for MI+ and MI− mice. A linear mixed-effects model (*R*^2^ = 0.27) with pellet speed (3-7 cm/s) and group (MI+ vs. MI−) as fixed effects and mouse (7 levels) as random effect indicated a significant main effect for pellet speed (*t*(700) = 9.289, *p* < 0.001) but no significant main effect for group or the interaction between group and pellet speed (all *p* > 0.05). Right: The same but for late learning. A linear mixed-effects model (*R*^2^ = 0.52) revealed significant main effects of pellet speed (*t*(839) = 3.408, *p* < 0.001), group (*t*(839) = −2.709, *p* = 0.007), and a significant interaction between pellet speed and group (*t*(839) = −3.647, *p* < 0.001). Data was first averaged across early or late days for individual mice and then across mice. Error bars denote the SEM across mice (N = 4 MI+ and 3 MI− mice). **(f)** Left: Lick speed for pellet speeds 3-7 cm/s during early learning for MI+ and MI− mice. A linear mixed-effects model (*R*^2^ = 0.16) with pellet speed and group as fixed effects and mouse as random effect indicated a significant main effect for pellet speed (*t*(700) = 3.95, *p* < 0.001), no significant main effect for group (*t*(700) = 1.149, *p* = 0.251), and a significant interaction between group and pellet speed (*t*(700) = −2.411, *p* = 0.016). Right: The same but for late learning. A linear mixed-effects model (*R*^2^ = 0.24) revealed significant main effects of pellet speed (*t*(839) = 4.74, *p* < 0.001), no significant main effect for group (*t*(839) = 1.078, *p* = 0.281), and a significant interaction between pellet speed and group (*t*(839) = −4.276, *p* < 0.001). Data was first averaged across early or late days for individual mice and then across mice. Error bars denote the SEM across mice (N = 4 MI+ and 3 MI− mice). **(g)** Left: Hit trial percentage for pellet speeds 3-7 cm/s during early learning for MI+ and MI−mice. A generalized mixed-effects model (*R*^2^ = 0.14) with pellet speed and group as fixed effects and mouse as random effect indicated a significant main effect for pellet speed (*t*(700) = −6.638, *p* < 0.001), a significant main effect for group (*t*(700 = −6.03, *p* < 0.001), and a significant interaction between group and speed (*t*(700) = 5.93, *p* < 0.001). Right: The same but for late learning. A linear mixed-effects model (*R*^2^ = 0.24) revealed significant main effects of pellet speed (*t*(839) = −5.353, *p* < 0.001), no significant main effect for group (*t*(839) = −1.602, *p* = 0.11), and a significant interaction between pellet speed and group (*t*(839) = 3.498, *p* < 0.001). Data was first averaged across early or late days for individual mice and then across mice. Error bars denote the SEM across mice (N = 4 MI+ and 3 MI− mice).

To better investigate motor preparation and execution, we divided each trial into segments based on the distribution of lick onsets: Approximately 99.99% of licks were initiated when the stage was within −3.4 (to the left) to 2.5 cm (to the right) of the midpoint of the mice. Based on this, we divided each trial into pre-, peri-, and post-target intervals (Figure 1c). At speeds 1-2 cm/s, the pre-target interval was 24-12 s and the peri-target interval was 5.9-3 s, whereas at speeds 3-7 cm/s these intervals ranged between 8-3.4 s and 2-0.84 s, respectively (Figure S1c). Moreover, the interquartile range of lick onsets in hit trials decreased compared to miss trials, suggesting that mice learned to initiate successful licks within 250-160 ms at speeds 1-2 cm/s, and within 130-60 ms at speeds 3-7 cm/s (Figure S1d). Since mice showed an increase in hit trial performance between the first and last three sessions only at speeds 3-7 cm/s (Figure S1b), the following analyses were mainly restricted to the pre- and peri-target intervals at those pellet speeds.

To assess learning-related changes in lick parameters, we calculated a modulation index (MI) for each mouse during early and late learning (Figure 1d). This index quantifies the degree to which mice modulate their first lick in anticipation of the approaching food pellet by combining three variables: (1) the absolute distance between the midpoint of the mouse and the pellet at the time of the lick (i.e., lick onset distance) with lower values indicating earlier licks, (2) the correlation between lick onset distance and pellet speed, with negative values indicating earlier licks for faster speeds, and (3) the correlation between lick speed and pellet speed, with positive values indicating faster licks for faster pellet speeds. We have previously shown that, early in learning, mice tend to lick faster for faster pellet speeds (3) but then shift towards initiating earlier licks (1) as well as licking earlier for faster pellet speeds (2) during late learning (Eltabbal et al., submitted).

Here, we identified two groups of mice during late learning that differed along this continuum of expected lick modulation (MI+ and MI−; Figure 1d). MI+ mice learned to initiate their first lick increasingly early and tended to lick slightly earlier with increasing pellet speed, whereas MI− mice learned to lick as the pellet approached their midpoint (Figure 1e). This pattern of licking was even more pronounced when considering successful trials during late learning only (Figure S1e, left panel). Figure S1f depicts all first lick onsets by an MI+ mouse for one example pellet speed, highlighting increasingly early licking over learning. Furthermore, MI− licked faster for faster pellet speeds during early and late learning, whereas MI+ mice showed a weaker relationship between lick speed and pellet speed, particularly during late learning (Figure 1f). This pattern tended to hold for successful trials during late learning (Figure S1e, right panel). While both groups improved their first lick performance, MI+ mice outperformed MI− mice on faster pellet speeds during late learning (Figure 1g). These findings validate our previous description of task-related learning, allowing us to investigate the contribution of MLIs to more anticipatory (MI+) and more reactive (MI−) licking.

### Lobule-specific Ca^2+^activity emerges with learning

We used one-photon widefield microscopy to monitor MLI Ca^2+^ activity dynamics by expressing GCaMP7f in the left cerebellar hemisphere via Cre-mediated conditional gene expression (Crus I and II: 5 c-kit/Cre mice; Crus I-only: 2 c-kit/Cre mice; Figure 2a). This region is part of a cortico-cerebellar loop involved in motor planning of orofacial movements, and MLI Ca^2+^ activity in Crus II has been previously associated with licking behavior (Astorga et al., 2017; Gaffield & Christie, 2017; Zhu et al., 2023). We employed *CellSort* to identify regions-of-interest (ROIs) based on ΔF/F0-normalized widefield recordings (Mukamel et al., 2009). Although one-photon imaging does not provide single-cell resolution of MLIs, *CellSort* allowed us to extract spatiotemporal fluctuations in the MLI response within each lobule and remove components overlying blood vessels (Crus I: 13.8 ± 0.8, Crus II: 16.5 ± 2 ROIs per session; mean ± SEM, Crus I: N = 7 mice, Crus II: N = 5 mice; Figures 2b and 2c). A similar PCA-ICA-based approach to identify ROIs and exclude hemodynamic activity has previously been employed by Makino et al. (2017) as well as Ren and Komiyama (2021). No bleaching on a per-trial basis was observed (Figure S2a).

**Figure 2:**
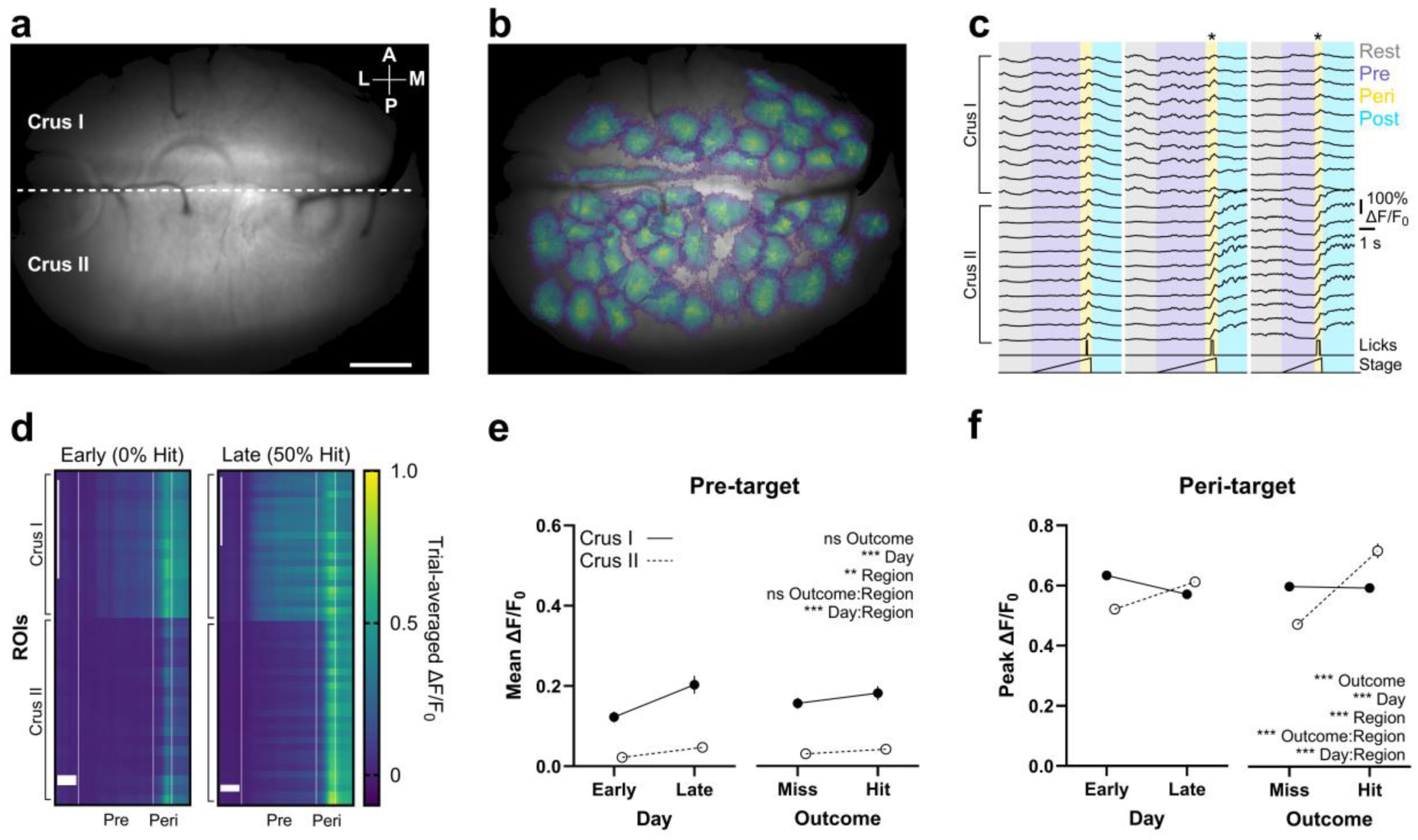
Wide-field imaging of interneurons reveals lobule-specific Ca^2+^ dynamics during learning. (a) Thresholded field-of-view for one example mouse. Scale bar = 500 μm. **(b)** Same field-of-view with overlaid regions-of-interest (ROIs) identified via CellSort (Mukamel et al., 2009). ROIs are color-coded according to weight, with yellow indicating the highest weight. **(c)** Example ΔF/F_0_ activity of a subset of ROIs during three example trials. Asterisks denote successful licks. **(d)** Trial-averaged ΔF/F_0_ activity over all trials of speed 6 cm/s within an early and a late session for the same mouse. ROIs in Crus I and II were sorted from lateral to medial. Vertical lines mark the start of the pre-, peri-, and post-target intervals. Vertical scale bars = 10 ROIs; horizontal scale bars = 1 s. **(e)** Line plots of the mean ΔF/F_0_ amplitude during the pre-target interval for Crus I and II grouped by day (early vs. late) and outcome (hit vs. miss). A linear mixed-effects model (*R*^2^ = 0.51) with mouse (7 levels) and pellet speed (5 levels) as random effects revealed significant main effects of day (*t*(2819) = 4.625, *p* < 0.001) and region (*t*(2819) = 2.67, *p* = 0.008) as well as a significant interaction between day and region (*t*(2819) = 8.902, *p* < 0.001), but no significant main effect of outcome or interaction between outcome and region (all other *p* > 0.05). **(f)** Line plots of the peak peri-target ΔF/F_0_ amplitude in Crus I and II grouped by day and outcome. A linear mixed-effects model (R^2^ = 0.35) revealed significant main effects of day (*t*(2819) = 5.21, *p* < 0.001), region (*t*(2819) = 19.467, *p* < 0.001), and outcome (*t*(2819) = 31.203, *p* < 0.001). The model also showed significant interactions between region and day (*t*(2819) = −11.683, *p* < 0.001) and outcome and region (*t*(2819) = −23.21, *p* < 0.001). Data plotted in **(e)** and **(f)** were averaged over trials within a session and then over mice. Error bars denote the SEM across mice (Crus I: N = 7 mice; Crus II: N = 5 mice).

Across learning, MLI activity in Crus I was higher than in Crus II during the pre-target interval (Figure 2d). Although we labeled and recorded a large field-of-view (FOV) over Crus I and II, the activity in individual ROIs within each lobule was low-dimensional (Figures 2c and 2d). Therefore, for all following analyses, we averaged the activity of all ROIs within Crus I and II on every trial. Crus I showed a larger increase from early to late learning compared to Crus II, but both regions were not affected by trial outcome (Figure 2e). During the peri-target interval, Crus II activity increased across learning, while activity in Crus I declined (Figure 2f). Crus II also increased in response to hit trials. Furthermore, at rest, Crus I activity decreased slightly from early to late learning whereas Crus II remained constant, suggesting that baseline amplitude differences cannot explain these opposing dynamics (Figure S2b). In addition, hit trials evoked rhythmic orofacial oscillations after lick offset (i.e., during the post-target interval), consistent with chewing-related movements (Barrett et al., 2024; Figure S2c). These movements were accompanied by a sustained increase in Crus II but not Crus I activity (Figure S2d). These findings suggest a lobule-specific division of labor: while Crus I becomes increasingly tuned to the motor preparation phase, Crus II shows pronounced activity during motor execution as well as pellet consumption.

### Persistent Ca^2+^ activity emerges together with anticipatory licking during learning

To investigate how MLI activity relates to motor preparation, we examined neural responses during the pre-target window. We only selected trials with no licks during this interval and at least one lick during the peri-target interval. Crus I activity increased from early to late learning and exhibited a plateau period - particularly in MI+ mice (Figure 3a). We therefore extracted the inflection point and duration of this plateau phase. Over the course of learning, MLI activity in MI+ mice ramped up earlier and remained elevated longer compared to MI− mice (Figures 3b and 3c). The inflection point remained stable on the distance-axis but became graded by speed on the time-axis, suggesting that persistent activity is distance-locked in both MI+ and MI− mice (Figure 3d). However, the prolonged plateau phase in MI+ mice became linked to earlier licking across days, whereas MI− mice showed a weaker relationship (Figure 3e). This finding suggests that increased persistent activity coincides with anticipatory licking in MI+ mice to a greater extent than in MI− mice. In addition, the inflection point of persistent activity is linked to the spatial position of the pellet.

**Figure 3:**
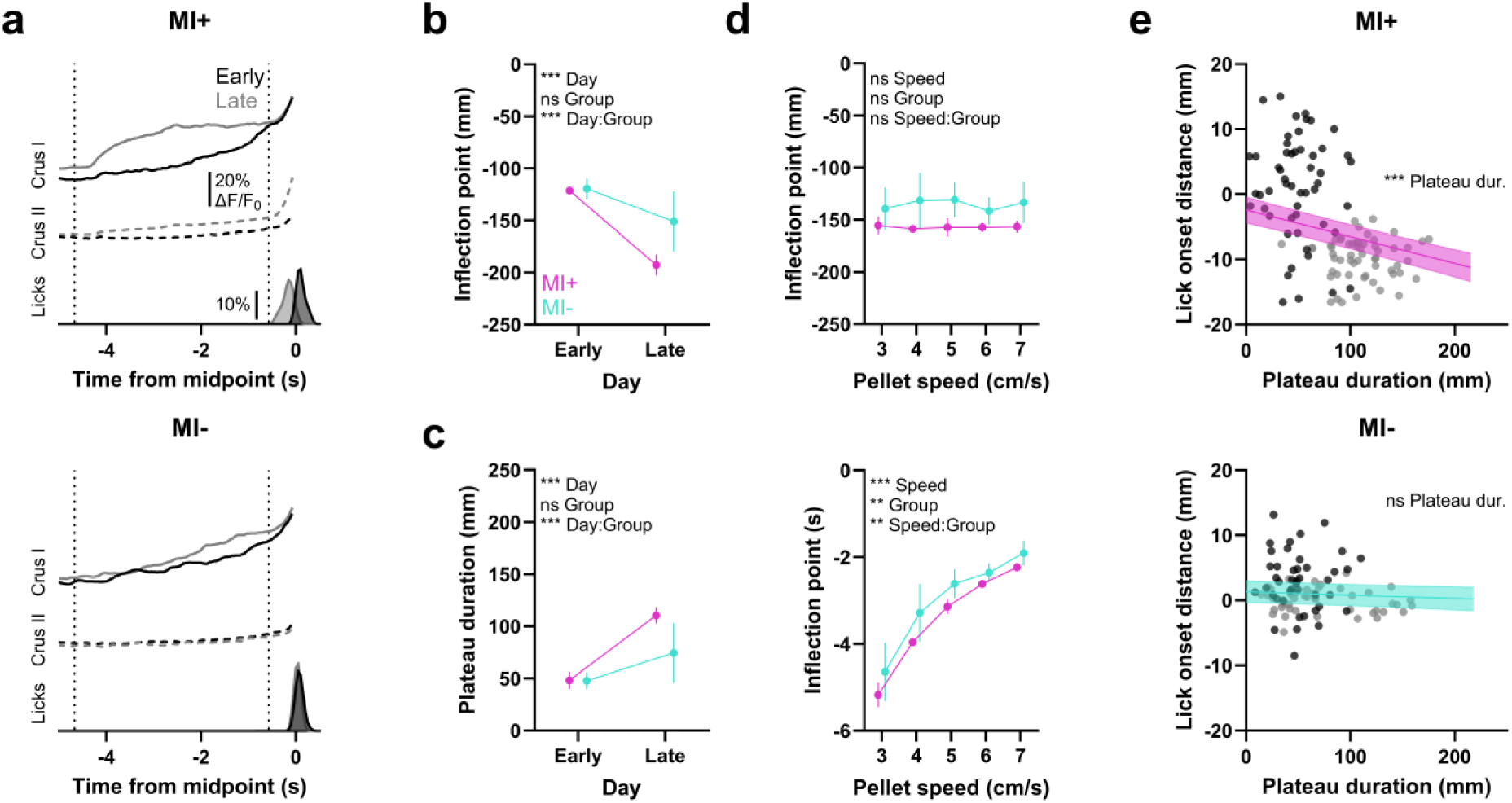
Persistent activity emerges with anticipatory licking in MI+ mice. (a) Representative mean ΔF/F_0_ traces in Crus I and II, along with lick probability histograms for an MI+ and MI− mouse. **(b)** Inflection point distance during early and late learning for MI+ and MI− mice. A linear mixed-effects model (*R*^2^ = 0.3) with group (MI+ vs. MI−) and day (early vs. late) as fixed effects and mouse (7 levels) and pellet speed (5 levels) as random effects showed a non-significant main effect for group (*t*(1536) = −0.23, *p* = 0.818), a significant main effect for day (*t*(1536) = −8.256, *p* < 0.001), and a significant interaction between group and day (*t*(1536) = −6.794, *p* < 0.001). Error bars denote the SEM over mice (N = 4 MI+ and 3 MI− mice). **(c)** Plateau duration during early and late learning for MI+ and MI− mice. A linear mixed-effects model (*R*^2^ = 0.34) with group and day as fixed effects and mouse and pellet speed as random effects showed a non-significant main effect for group (*t*(1536) = 0.219, *p* = 0.827), a significant main effect for day (*t*(1536) = 9.016, *p* < 0.001), and a significant interaction between group and day (*t*(1536) = 7.237, *p* < 0.001). Error bars denote the SEM over mice (N = 4 MI+ and 3 MI− mice). **(d)** Top: Inflection points on the distance-axis for speeds 3-7 cm/s across learning. A linear mixed-effects model (*R*^2^ = 0.13) with pellet speed and group as fixed effects and mouse as random effect showed no significant main effects for pellet speed and group, nor a significant interaction (all *p* > 0.05). Bottom: Inflection points on the time-axis. A linear mixed-effects model (*R*^2^ = 0.37) revealed significant main effects for pellet speed (*t*(2116) = 19.824, *p* < 0.001) and group (*t*(2116) = −2.308, *p* = 0.021), and a non-significant interaction between pellet speed and group (*t*(2116) = 1.419, *p* = 0.156). Error bars denote the SEM over mice (N = 4 MI+ and 3 MI− mice). **(e)** Top: Lick onset distance as a function of plateau duration on the distance-axis for MI+ mice. A linear mixed-effects model (*R*^2^ = 0.09) with plateau duration as a fixed effect and mouse (4 levels) and pellet speed as random effects showed a significant main effect for plateau duration (*t*(1151) = −9.053, *p* < 0.001, Cohen’s *d* = 0.53). Bottom: Lick onset distance versus distance above threshold for MI− mice. A linear mixed-effects model (*R*^2^ = 0.16) with mouse (3 levels) and pellet speed as random effects revealed a non-significant main effect for plateau duration (*t*(965) = −1.92, *p* = 0.055, Cohen’s *d* = 0.12). Superimposed lines represent fitted regression lines from the trial-based linear mixed-effects models, with the shaded area indicating the 95% confidence intervals. Black and gray data points represent trial averages over the same speed within a session for individual mice during early and late learning, respectively.

Could factors such as motivation or increased responsiveness to reward explain the increase in persistent activity during late learning in MI+ mice? To address this, we examined the effect of trial sequence and outcome within a session on persistent activity. As mice completed more trials, the duration of persistent activity tended to decline (*t*(800) = −1.662, *p* = 0.097), but this trend did not differ between early and late learning (*t*(800) = 0.612, *p* = 0.541). Furthermore, hit trials exhibited a non-significant association with increased persistent activity (*t*(800) = 1.132, *p* = 0.258), but neither did this effect differ between early and late learning (*t*(800) = 0.449, *p* = 0.654; linear mixed-effects (LME) model; *R*^2^ = 0.33). These results suggest that factors related to session duration and reward responsiveness did not differentially affect MI+ mice during late learning.

Since the inflection point of persistent activity is distance-locked, we examined whether MLI activity is graded according to the speed of the approaching pellet. To avoid a spurious relationship due to time compression, we first identified the offset of the initial rise phase (i.e., onset of the plateau phase) and then calculated the mean derivative of the subsequent plateau phase on the time axis (Figure S3a). During early learning, the plateau derivative was inversely related to pellet speed, though less pronounced in MI+ mice (Figure S3b, left panel). During late learning, the plateau derivative increased with increasing pellet speed in both MI+ and MI−mice, suggesting that the rate of change of MLI activity increased with corresponding increases in pellet speed (Figure S3b, right panel). Therefore, we examined whether the variability in lick onsets due to pellet speed can be explained by the plateau derivative. Only in MI+ mice was earlier licking tied to a plateau derivative increase during late learning (Figure S3c, left panel). In addition, neither MI+ nor MI− mice exhibited a relationship between the plateau derivative and lick speed (Figure S3c, right panel). These results suggest that the rate of change of persistent activity during late learning is graded by the speed of the approaching pellet, which becomes associated with earlier licking for faster pellet speeds in MI+ mice, whereas MI− mice do not associate MLI dynamics with lick parameters.

### MI+ mice decouple lick initiation from immediate sensory input

To examine how lobule-specific dynamics relate to motor execution in MI+ and MI−mice, we examined neural responses during the peri-target window. For this analysis, we only used mice for which we imaged Crus I and II simultaneously and extracted the first lick onset in the peri-target interval that was separated by at least three seconds from any previous licks. Both groups exhibited a stable response in Crus I when the stage reached the midpoint of the mice, whereas Crus II responses shifted with lick onset in MI+ but not MI− mice (Figure 4a). If Crus I encodes a sensory signal at the midpoint, we hypothesized that aligning trials to the midpoint would reveal no differences between late-timed lick and no-lick trials, but that aligning to lick onset would show pronounced differences between early- and late-timed lick trials. Indeed, we observed that Crus I showed a similar response irrespective of licking, and this response was thus higher in amplitude during trials in which mice licked late (Figure 4b). Conversely, if Crus II reflects a motor response, we expected to find differences between late-timed lick and no-lick trials when aligned to the midpoint, but no difference between early- and late-timed lick trials aligned to lick onset (Figure 4b). Accordingly, even when the first lick was successful, the response in Crus I at lick onset decreased over time in MI+ but not MI− mice, whereas no difference was observed at lick onset for Crus II (Figure 4c). These findings suggest that activity in Crus I during the peri-target interval is temporally consistent over learning and does not adapt even when MI+ mice learn to lick earlier. Therefore, MLIs in Crus I may encode a stable sensory-related signal around the midpoint, such as a visual or proprioceptive response due to the passing of the stage, whereas MLI responses in Crus II are tied to lick onset and subsequent modulation.

**Figure 4:**
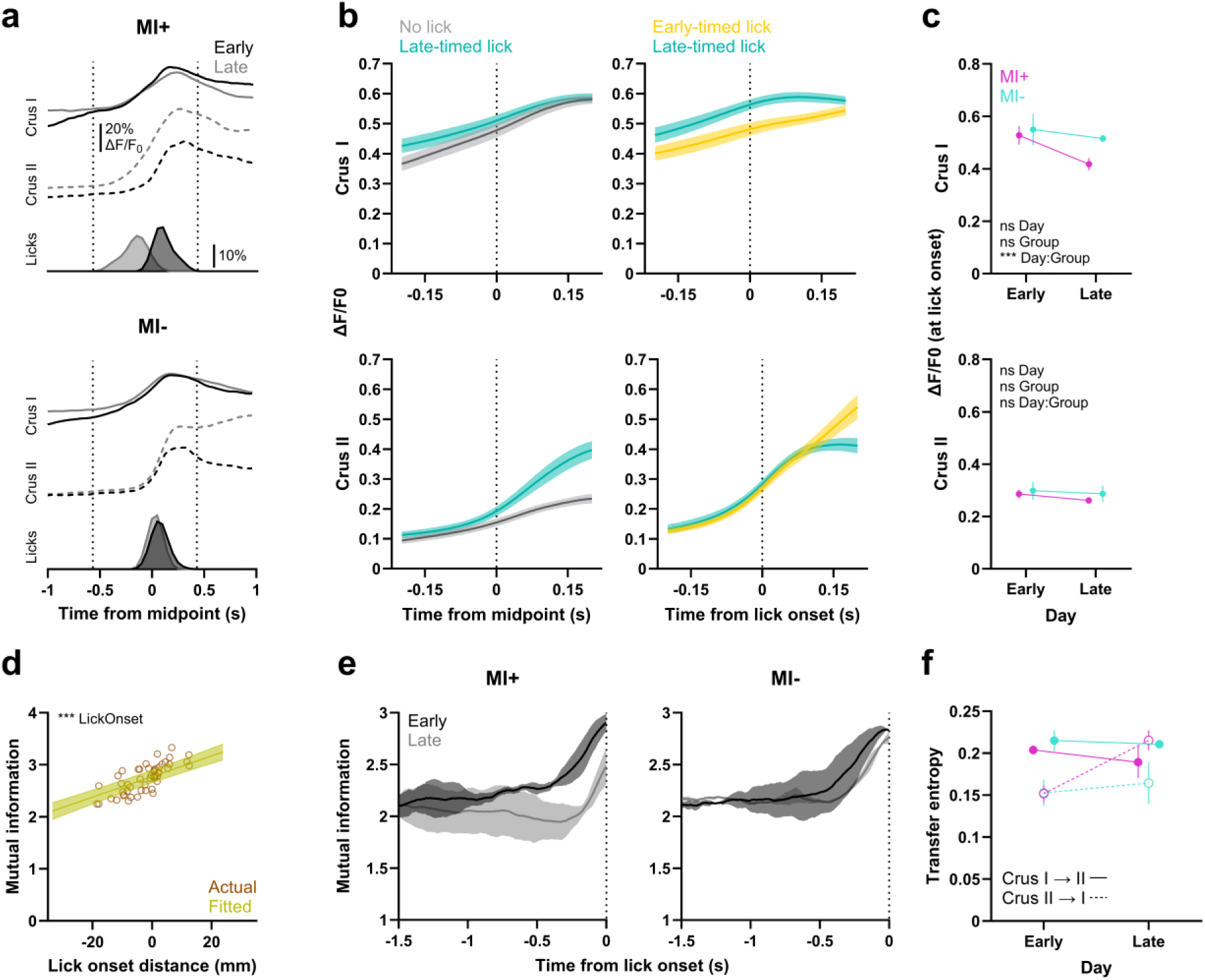
MI+ mice decouple licks from immediate sensory-related signals at the midpoint. **(a)** Representative mean ΔF/F_0_ traces in Crus I and II along with lick probability histograms for an MI+ and MI− mouse. **(b)** Left: ΔF/F_0_ in Crus I and II for trials with no licks and licks initiated after the midpoint of the mice (late-timed licks). Right: ΔF/F_0_ in Crus I and II for trials with licks initiated before the midpoint (early-timed licks) and late-timed licks. Data was first averaged over trials and then over mice. Error bars represent the SEM (N = 5 mice). **(c)** Top: Mean ΔF/F_0_ in Crus I in a 100 ms window centered around the lick onset in single-lick hit trials for MI+ and MI− mice. A linear mixed-effects model (*R*^2^ = 0.49) with group (MI+ vs. MI−) and day (early vs. late) as fixed effects and mouse (5 levels) and pellet speed (5 levels) as random effects showed non-significant main effects for group and day (all *p* > 0.05) and a significant interaction between group and day (*t*(353) = −4.62, *p* < 0.001). Bottom: Mean ΔF/F0 in Crus II around the lick onset in single-lick hit trials. A linear-mixed effects model (*R*^2^ = 0.21) showed no significant main effect for group and day or their interaction (all *p* > 0.05). Data was first averaged over trials and then over mice. Error bars represent the SEM (N = 3 MI+ and 2 MI− mice). **(d)** Mutual information between Crus I and II in a 500 ms window before lick onset. A linear mixed-effects model (*R*^2^ = 0.19) with lick onset as fixed effect and mouse and pellet speed as random effects showed a significant main effect of lick onset (*t*(1062) = 11.177, *p* < 0.001). Data points represent trial averages over the same speed within early or late learning for individual mice. The superimposed line represents the fitted regression line from the trial-based linear mixed-effects model, with the shaded area indicating the 95% confidence interval (N = 5 mice). **(e)** Mutual information between Crus I and II calculated in 500 ms moving windows until lick onset for MI+ and MI− mice. A linear mixed-effects model (*R*^2^ = 0.17) to explain mutual information in the last 500 ms before lick onset with group and day as fixed effects and mouse and pellet speed as random effects showed non-significant main effects for group and day (all *p* > 0.05) and a significant interaction between group and day (*t*(1060) = −5.803, *p* < 0.001). Data was first averaged over trials and then over mice. Error bars represent the SEM (N = 3 MI+ and 2 MI− mice). **(f)** Transfer entropy between Crus I and Crus II during early and late learning for MI+ and MI−mice. A linear mixed-effects model (*R*^2^ = 0.18) to explain transfer entropy from Crus I to Crus II with group and day as fixed effects and mouse and pellet speed as random effects revealed no significant main effects (all *p* > 0.05) and a significant interaction between group and day (*t*(1060) = −3.903, *p* < 0.001). A second linear-mixed-effects model (*R*^2^ = 0.28) for the reverse direction (Crus II to Crus I) showed a non-significant main effect of group (*t*(1060) = −0.271, *p* = 0.786), a significant main effect of day (*t*(1060) = 2.673, *p* = 0.008), and a significant interaction between group and day (*t*(1060) = 8.905, *p* < 0.001). Data was first averaged over trials and then over mice. Error bars represent the SEM (N = 3 MI+ and 2 MI− mice).

To assess whether learning alters the degree and direction of interactions between Crus I and II, we analyzed mutual information and transfer entropy. A similar approach has been used to demonstrate altered coupling between regions involved in movement planning and execution (Lizier et al., 2011). Importantly, we do not assume inter-lobular connectivity but rather intended to test whether local or input processing in Crus I and II becomes functionally decoupled during learning. Across all mice, mutual information increased together with lick onset (Figure 4d). However, in MI+ mice, mutual information declined over learning approximately 500 ms before lick onset (Figure 4e). In addition, the direction of information flow reversed in MI+ but not MI− mice: Transfer entropy from Crus I to Crus II decreased, and it increased from Crus II to Crus I over learning. MI− mice only exhibited a comparably small increase in transfer entropy from Crus II to Crus I (Figure 4f). We chose a delay parameter of 200 ms for calculating transfer entropy and confirmed that changing the delay parameter between 100 to 500 ms did not change the direction and statistical significance of the effects (data not shown). These results suggest that MI+ mice learn to decouple motor responses from immediate sensory-related signals over learning.

We hypothesized that the dynamics of the midpoint signal carry information regarding lick parameters specifically in MI− mice. To test this idea, we extracted the mean derivative of the Crus I Ca^2+^ transient during 500 ms before lick onset (Figure S4a). The derivative in Crus I was positively graded by pellet speed, but, consistent with decoupling, this graded relationship was less pronounced in MI+ mice during late learning (Figure S4b). Early in learning, faster licks were already associated with a higher derivative (*t*(698) = 3.806, *p* < 0.001; main effect of derivative) but less pronounced in MI+ mice (*t*(698) = −2.481, *p* = 0.013; interaction between group and derivative; LME; *R*^2^ = 0.16; all other *p* > 0.05). This relationship was particularly evident during late learning (Figure S4c). We separately confirmed that the derivative in Crus II was not related to lick speed during early (*t*(480) = 0.681, *p* = 0.496; main effect of derivative; LME; *R*^2^ = 0.12; all other *p* > 0.05) or late learning (*t*(574) = 1.935, *p* = 0.054; main effect of derivative), and this relationship even tended to reverse for MI+ mice (*t*(574) = −2.102, *p* = 0.036; interaction between group and derivative; LME; *R*^2^ = 0.23). Furthermore, higher mean derivatives were related to later lick onset during early learning (*t*(700) = 5.259, *p* < 0.001; main effect of derivative), but this relationship was more pronounced in MI+ mice (*t*(700) = 2.907, *p* = 0.004; interaction between group and derivative; LME; *R*^2^ = 0.22; all other *p* > 0.05). We observed the same pattern of effects during late learning (Figure S4c). These results suggest that the midpoint signal in Crus I does not dynamically adjust to account for earlier licking. Moreover, MI+ mice dissociate lick dynamics, whereas MI− mice tune lick speed in line with immediate sensory-related signals as the pellet approaches the midpoint.

### Persistent activity is tied to the spatial position of sensory cues

We used generalized linear models to examine the unique contributions of the pellet’s spatial position, body kinematics, licks, and reinforcement to MLI activity during the pre- and peri-target intervals (Figures 5a, 5b and 5c). Since we previously found that mice show seconds-long preparatory body movements (El-Tabbal et al., submitted), we specifically aimed to test whether MLI activity simply reflects body movement. We therefore included several uncorrelated body movement parameters and hypothesized pellet position, but not body kinematics, to become an important predictor during the pre-target interval, particularly for MI+ mice. As expected, β coefficients for pellet position increased from early to late learning, and this increase was pronounced in Crus I and in MI+ mice (Figure 5d). β coefficients for body kinematics were also higher in Crus I than Crus II, but decreased over time, especially in Crus I, and MI+ mice tended to show a more pronounced decrease in β coefficients though not statistically significant. Licks and reinforcement showed little contribution to activity during the pre-target interval. These results suggest that the spatial position of the pellet becomes increasingly linked to MLI activity, particularly in mice that develop anticipatory lick responses.

**Figure 5:**
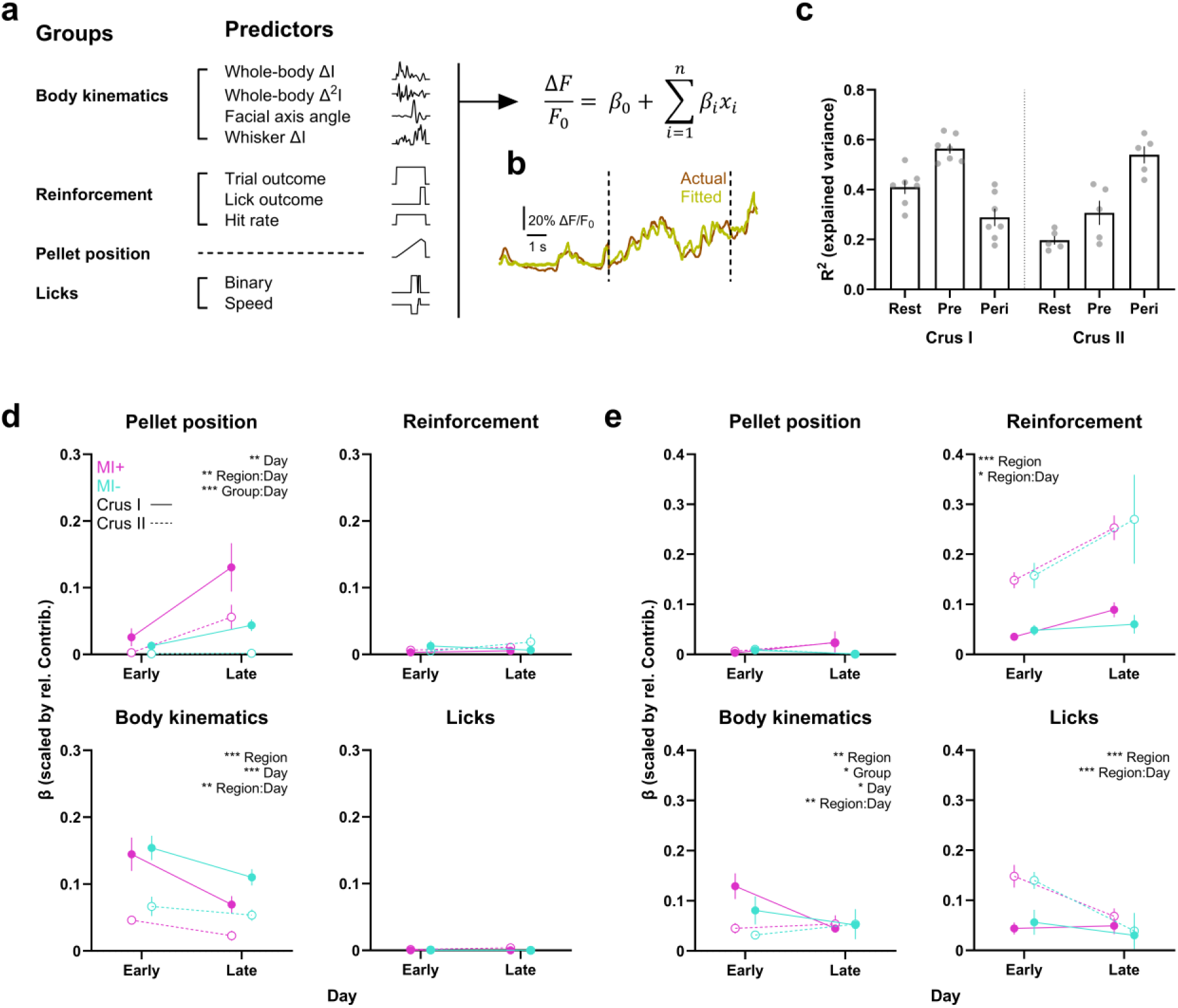
Interneuron activity during the pre-target interval becomes linked to pellet position in MI+ mice. **(a)** Continuous, whole-trial, and event variables associated with each predictor group along with the generalized linear model formula. ΔI = intensity rate; Δ^2^I = intensity acceleration. **(b)** Actual and fitted traces for one example trial. **(c)** Explained variance for each trial interval for Crus I and Crus II. *R*^2^ was computed for each day, averaged over days, and then averaged over mice. Error bars denote the SEM (Crus I: N = 7 mice; Crus II: N = 5 mice). **(d)** Scaled β weights for each predictor group during the pre-target interval divided by MI+ and MI− mice as well as Crus I and II. For pellet position, a linear mixed-effects model (*R*^2^ = 0.74) with the factors region (Crus I vs. Crus II), group (MI− vs. MI+), day (early vs. late), and their two-way interactions revealed a significant main effect of day (*t*(65) = 2.882, *p* = 0.005), a significant interaction between region and day (*t*(65) = −2.953, *p* = 0.004), and a significant interaction between group and day (*t*(65) = 4.497, *p* < 0.001). For body kinematics, a mixed-effects model (*R*^2^ = 0.76) revealed significant main effects for region (*t*(65) = −7.302, *p* < 0.001) and day (*t*(65) = 4.15, *p* < 0.001), and a significant interaction between region and day (*t*(65) = 3.156, *p* = 0.002). The interaction between group and day did not reach significance (*t*(65) = −1.655, *p* = 0.103). Linear mixed-effects models for reinforcement and licks did not reveal any significant effects during the pre-target interval (all *p* > 0.05). Data shown was first averaged over early or late days for individual mice and then across mice. Error bars represent the SEM (Crus I: N = 4 MI+ and 3 MI− mice; Crus II: N = 3 MI+ and 2 MI− mice). **(e)** Same as in **(d)** but for the peri-target interval. A linear mixed-effects model for pellet position (*R*^2^ = 0.18) did not reveal any statistically significant effects, though the interaction between group and day (*t*(65) = 1.798, *p* = 0.077) showed the expected effect direction. For body kinematics, a mixed-effects model (*R*^2^ = 0.33) revealed significant main effects for region (*t*(65) = −3.023, *p* = 0.004) and day (*t*(65) = −2.056, *p* = 0.044), and a significant interaction between region and day (*t*(65) = 3.443, *p* = 0.001). The interaction between group and day did not reach significance (*t*(65) = −1.724, *p* = 0.09). A mixed-effects model for reinforcement (*R*^2^ = 0.57) showed significant main effects for region (*t*(65) = 3.751, *p* < 0.001), and a significant interaction between region and day (*t*(65) = 2.002, *p* = 0.049). For licks, mixed-effects modeling (*R*^2^ = 0.577) revealed significant main effects of region (*t*(65) = 5.425, *p* < 0.001), and a significant interaction between region and day (*t*(65) = −4.184, *p* < 0.001).

During the peri-target window, we expected licks and reinforcement to be represented in Crus II, and the contribution of pellet position to increase in MI+ compared to MI− mice late in learning due to sensorimotor decoupling. Reinforcement elicited larger β coefficients in Crus II than Crus I, and reinforcement tended to become more important over the course of the experiment (Figure 5e). Licks also elicited larger β coefficients in Crus II than Crus I, but licks became less important for explaining overall ΔF/F0 activity during late learning. In addition, MI+ mice showed an increase in the contribution of pellet position over time, but this effect did not reach statistical significance. These results confirm that licks and reinforcement are more strongly represented in Crus II.

## Discussion

To investigate how MLIs contribute to dynamic sensorimotor control, we used a task in which mice freely initiated licks to intercept moving food pellets. We observed that MLIs in Crus I developed persistent Ca^2+^ activity over several seconds preceding licks in mice that exhibited more anticipatory behavior. This persistent activity was sensory-related and, with learning, encoded target motion independent of body movement. In addition, MLIs produced stable responses when the stage holding the food pellet passed immediately in front of the animal irrespective of licking, and mice that developed more anticipatory behavior thus learned to decouple motor responses from this midpoint signal. While previous work highlighted that MLI activity is related to two-dimensional stimulus-valence (Ma et al., 2020) or movement rate, but not to intentional motor adjustments (Gaffield & Christie, 2017), our findings indicate that MLIs in Crus I relay acquired sensory representations for the anticipatory control of tongue movements.

We observed persistent activity lasting several seconds consistent with recent reports of motor preparatory signals in the cerebellar hemispheres and sustained ramping activity in granule cells (Gao, 2018; Garcia-Garcia, 2024; Prat et al., 2024; Zhu et al., 2023). For example, Gao (2018) demonstrated that the cerebellar nuclei exhibit persistent activity driven by frontal cortex up to 1.3 seconds before movement onset. Recently, Garcia-Garcia et al. (2024) found that granule cells display ramping activity that aligns with delay periods of up to 2 seconds preceding an expected reward. These forms of persistent activity have been interpreted as signals of urgency or reward expectation, respectively. If interneurons in our task were encoding expectation or urgency, we would expect MLI activity to ramp up later for slower pellet speeds without a relationship with velocity during the plateau period. However, persistent activity during the pre-target interval was related to pellet position independent of body movement, and MLIs produced a stable response when the pellet reached the midpoint irrespective of licking, which supports the interpretation that MLIs in Crus I encode sensory representations. We can only speculate that MLI activity is driven by granule cells and may therefore contribute to excitation-inhibition balance of Purkinje cells (Jelitai et al., 2016), though cerebellar nucleo-cortical mossy fibers (Gao et al., 2016) as well as spillover glutamate from climbing fibers (Szapiro & Barbour, 2007) are alternative hypotheses. Finally, future studies may uncover to what extent the increased MLI activity in Crus I observed here reflects distinct cognitive components, such as visuospatial attention, mediated via connectivity between the cerebellum and prefrontal cortex (Squire et al., 2013).

Previous research has emphasized the contribution of sensory processing within the cerebellar hemispheres. In humans, the dentate nuclei show activity during sensory stimulation rather than motor control, suggesting that the lateral cerebellum regulates sensory data acquisition (Baumann et al., 2015; Bower, 1997; Gao et al., 1996). In addition, Crus I was shown to be involved in visual processing of biological motion and to increase its activity with perceptual task demands (Baumann et al., 2010; Sokolov et al., 2012). In rodents, Michikawa et al. (2021) demonstrated that Crus I along with vermal lobules V-VIII show increased complex spike activity in response to somatosensory stimulation compared with other cerebellar lobules. MLIs in Crus I have also recently been shown to exhibit sensory-driven (i.e., puff-evoked) synchrony that affects subsequent whisker protractions (Brown et al., 2025). Our GLM analysis during the pre-target interval suggests that MLI activity in Crus I initially reflects body movement but, over the course of learning, is uniquely explained by pellet position. However, while our wide-field imaging captured a mixed signal likely reflecting stellate cell activity, we cannot distinguish between stellate and basket cells as well as the recently defined MLI subtypes 1 and 2 (Kozareva et al., 2021). Future subtype-specific analyses may reveal whether MLI1s maintain persistent activity during sensory processing, while MLI2s gate its timing or modulate its amplitude (Lackey et al., 2024; Park et al., 2026). Further investigating how MLI activity in Crus I may capture statistics of the environment and thereby affect downstream processing represents an exciting research direction (Michikawa et al., 2021).

To what degree may eye movements explain persistent activity of MLIs? Crus I and II have recently been implicated in sensory data acquisition irrespective of pupil size and position (Khilkevich et al., 2024). In addition, eye-movements primarily elicit activity in the cerebellar vermis, flocculus, paraflocculus and the fastigial nuclei (Kheradmand & Zee, 2011). Cerminara et al. (2009) also demonstrated that Purkinje cells in Crus I exhibit elevated simple spike discharge in response to a moving visual target during an interception task. This tonic elevation was not correlated to eye or limb movements and persisted during temporary occlusion of the target and was therefore interpreted as an internal model of target motion (Cerminara et al., 2009). However, since we did not track pupil position, we cannot definitively exclude the influence of eye movements on the persistent activity we observed.

The mutual information and transfer entropy analyses indicate that, with learning, sensory and motor processing undergo temporal decoupling. The reactive responses (i.e., tuning lick speed with pellet speed) can be explained by the midpoint signal in Crus I whereas the anticipatory responses (i.e., tuning lick onset with pellet speed) are better aligned with the pre-target activity in Crus I. While conceptually similar to eyelid conditioning, our results point toward a broader framework of sensorimotor decoupling for continuous stimuli and motor responses. Recently, Boven et al. (2023) suggested a computational framework in which cerebro-cerebellar loops guide learning through feedback decoupling. In their model, future feedback signals are predicted by the cerebellum and then relayed to the cerebral cortex, thereby reducing dependency on external feedback. This decoupling mechanism explains our result that, with learning, the sensory midpoint signal becomes separated from motor outputs in MI+ but not MI− mice. In addition, Boven et al. (2023) modeled the cerebellum as a forward model that learns to predict the sensory consequences of motor commands (Ito, 1970; Miall et al., 1993). Bilateral inactivation may reveal whether MLIs in Crus II relay an efference copy used for forward modeling by cerebellar circuits (Gaffield & Christie, 2017). Future studies may also test whether MLIs are involved in fine-graded control of the tongue, such as during acceleration and deceleration (Hage et al., 2025) or directional movement control in response to target motion (Bina et al., 2025).

## Methods

### Animals and surgery

All experimental procedures involving the use of animals have been approved by the Animal Care and Use Committee of the Okinawa Institute of Science and Technology (OIST). Surgeries were performed in an Association for Assessment and Accreditation of Laboratory and Animal Care (AAALAC International)-accredited facility. 28 c-kit/Cre mice (JAX #032923, The Jackson Laboratory; Amat et al., 2017) aged 8-12 weeks underwent chronic cranial window surgery over the left cerebellar hemisphere following our laboratory’s protocol (Augustinaite & Kuhn, 2020). Care was taken to ensure that the dura mater remained fully intact during the craniotomy. A 5 mm glass cover slip with three silicone access ports (Roome & Kuhn, 2014; 2019) was centered approximately 100-300 um posterior from the posterior suture line and 2.5-2.75 mm lateral and glued to the skull. Access ports were positioned along the border between Crus I and II based on the visible horizontal fissure to allow for virus injections and imaging in both lobules. All mice were monitored daily and given subcutaneous analgesia and saline injections for up to 4 days post-surgery. Mice were single housed in environmentally enriched cages (running wheel, dome, tunnel, and bedding). Handling and virus injections were initiated once mice recovered around 90 percent of their pre-surgery weight.

### Wide-field microscope setup

Wide-field calcium imaging was performed via one-photon excitation of fluorescence at 488 nm using a continuous wave laser (Genesis CX, Coherent) under a BOB microscope (Sutter Instrument) combined with a x5/N.A. 0.13 air objective (Zeiss). The laser was coupled to the microscope using a liquid light guide, and the light was collimated via the built-in Olympus vertical illuminator. We used an excitation filter (470/40), a dichroic mirror (495), and an emission filter (525/50). Recordings were made using a CMOS camera (ATLAS 10, LUCID Vision Labs, Inc.) together with Arena SDK software (LUCID Vision Labs, Inc.). Frames were acquired at 30.7 Hz across a field-of-view of 3.28 x 3.92 mm.

### Virus injections

Mice were head-mounted on the wide-field microscope setup and anesthetized with isoflurane (1-2%). Beveled quartz pipettes (10 µm tip diameter) angled at 30° were used to inject viral solution through the silicone access port with the tip flange oriented towards the brain surface. For Ca^2+^ imaging of MLIs, each mouse received between 3-5 and 5-6 50 nl injections of pGP-AAV9-syn-FLEX-jGCaMP7f-WPRE (titer: 2.8 x 10E13; gifted by Douglas Kim and GENIE Project; Addgene viral prep # 104492-AAV9; Dana et al., 2019) in Crus I and II, respectively. Injections were performed at low pressure (PSI < 0.3) at 50 µm depth to target the molecular layer. The pipette was retracted after waiting for five minutes. A drop of purified water was applied to the tip of the pipette after every injection to unclog the pipette.

### Behavioral training

A 10-day training protocol was initiated after mice recovered from surgery. At the start of each day, mice were transferred to the experiment room and allowed to acclimatize for 30 minutes. Mice were tunnel-handled repeatedly during 5-minute sessions on the first three days. Virus injections were made on day 3. Afterwards, mice were put on water restriction for four days followed by food restriction which lasted until the end of the experiment. During water restriction, mice were guided into a modified 3D-printed body restraint tube (Wagner et al., 2020) and head-fixed to two horizontal head bars. Habituation to head-fixation lasted 15 minutes every day and mice were allowed to consume up to 1 ml water from a spout. Around 1 g water gel was added to each cage after returning the mice. During food restriction, mice were head-fixed and allowed to consume up to 0.5 g of the same food pellets (5TUL, 20 mg, TestDiet) that were used during the main experiment. The food pellets were individually positioned on a stage that moved towards the mice at low speed (< 0.5 cm/s) and stopped directly in front of the mouse. On the last day of training, mice completed an experimental session comprising approximately 40 counterbalanced and randomized speeds of 1 and 2 cm/s. All mice were maintained at around 80-90 percent of their post-surgery weight by adding 1-2 g of food pellets to their cages. Weights were monitored daily before the start of each session.

### Behavioral setup

The behavioral setup was previously described in detail by Eltabbal et al. (submitted). We used a belt-driven linear actuator (W45-15, CCM Automation Technology Co., Ltd.) together with a servo motor (ClearPath-SDSK, Teknic, Inc.). The motor was controlled using the ClearCore library (Teknic, Inc.) via an Arduino microcontroller (Mega 2560). The platform holding the food pellet was 3D-printed and attached to a micromanipulator (M-3333, Narishige Co., Ltd.) that was placed on the linear actuator. We used a pellet dispenser (PD-020D, O’Hara Co., Ltd.) to dispense pellets automatically at the start of each trial if a photosensor (ST0250, SunFounder) detected light from a laser (KY008-650 nm) due to the absence of a pellet on the platform. One blue LED illuminated the setup throughout the experiments (M470L5, Thorlabs, Inc.). Stage movement and pellet refilling were automatically controlled using custom-written scripts in C++ using the Arduino integrated development environment.

### Targeted lick-interception task

For the functional imaging experiment, seven mice completed eight consecutive daily sessions of the targeted lick-interception task. During each trial, a single food pellet was dispensed onto the motorized stage that moved toward the animal at one of seven discrete speeds (1-7 cm/s), traveling from a position 27.5 cm to the left of the animal to a position 2.5 cm to the right. To distinguish between the pellet moving toward and away from the mouse, the distance between the mouse and pellet ranged from −27.5 to 2.5 cm. Each session lasted 1-1.5 hours and typically contained around 70 randomized and counterbalanced trials (ten trials per speed). Up to ten seconds of spontaneous rest activity was recorded before the start of every trial, which was signaled with a 500 ms tone. The noise level at the experimental set-up was measured at 65.5 decibels (dB). Assuming a just-noticeable-difference of 1 dB, white noise was played at 74.4 dB to mask any sounds generated by the motor. The inter-trial interval was randomly selected between 15 and 25 seconds. We defined the peri-target window as the four-sigma (± 4σ, where σ = 1.48 x mean absolute deviation) range around the mean lick onset distance across all mice, pellet speeds, and days.

### Modulation index

To investigate learning-related changes in lick parameters, we calculated a modulation index (MI) as described in Eltabbal et al. (submitted):

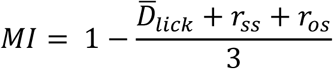

where 𝐷̅_𝑙𝑖𝑐𝑘_denotes the mean lick onset distance, 𝑟_𝑜𝑠_denotes the Pearson correlation between the lick onset distance and pellet speed, and 𝑟_𝑠𝑠_denotes the Pearson correlation between the lick speed and pellet speed. We calculated these variables via bootstrapping by averaging over 1000 iterations in which one trial per pellet speed was selected and the average lick onset distance and correlations were calculated. 𝐷̅_𝑙𝑖𝑐𝑘_was normalized across all mice to a range between 0 and 1. The correlation coefficients were normalized to fall within a range between 0 and 1.

### Movement tracking

Two 500 Hz infrared cameras (Blackfly S USB3, Teledyne FLIR LCC) were positioned laterally (side-view) and ventrally (bottom-view) to capture mouse behavior. Four infrared LEDs (M940L3, M680L5, Thorlabs, Inc.) were placed alongside. Videos were recorded using Spinnaker SDK (Teledyne Vision Solutions). Separate *DeepLabCut* (Mathis et al., 2018; Nath et al., 2019) networks using the Resnet50 model were trained repeatedly for 200,000 iterations on a subset of frames manually annotated for the tongue, paws, and snout. The trained *DeepLabCut* models were applied to all behavioral videos, and the resulting coordinates were filtered with a 95 percent likelihood threshold. Tongue, paw, and snout coordinates were further combined into three-dimensional representations from which speed, acceleration and distance were determined. Whisker pad, right eye, and whole-body motion were derived from manual masks for which the first or second derivative of the mean pixel intensity was calculated, reflecting changes in intensity rate or intensity acceleration.

### Motion correction

We used FlowReg (Flotho et al., 2022) implemented in MATLAB (MathWorks, Inc.) to correct rigid and non-rigid displacements within the field-of-view for all recordings within a session. Corrected recordings from the same mouse were then registered to a common reference template across all days using the *imregtform* function as described by Makino et al. (2017).

### Extraction and normalization of Ca^2+^ signals

We used CellSort (Mukamel et al., 2009) to extract independent components (ICs) from MLI recordings via temporal principal component analysis (PCA) and spatiotemporal independent component analysis (ICA). All recordings were first thresholded above the 40th percentile of the mean pixel amplitude across all sessions. Temporal PCA was then run on all frames recorded during stage movement within a single session. Principal components whose explained variance exceeded twice the estimated noise floor were retained for spatial ICA. A minimum threshold of 5000 pixels was used to filter out ICs close to the edges of the FOV. We calculated the z-scores of the weighted masks, retaining only pixels exceeding 1.5 standard deviations (SDs). All ICs were visually inspected, and those overlapping with blood vessels or the edges of the access ports were excluded from the analysis. The remaining ICs were normalized to sum to one and back-multiplied with the motion-registered recordings. The products for all pixels were added together to obtain weighted average traces for each ROI. Baseline fluorescence (F0) was estimated by averaging all frames recorded one second before or after any body movement during up to 10 seconds of rest pre-trial activity. We then normalized Ca^2+^ activity by subtracting F0 from the fluorescence at each frame and dividing by F0 to calculate ΔF/F0. These resulting traces were averaged across ICs within Crus I and Crus II for all analyses.

### Spectral analysis

We used the Fast Fourier Transform algorithm via MATLAB’s *fft* function to compute the single-sided amplitude spectrum of each trial’s whisker pad intensity rate. Frequencies were restricted between 0 to 20 Hz.

### Quantification of persistent activity

ΔF/F_0_ traces were first smoothed with a second-order 1 Hz low-pass Butterworth filter. The inflection point of persistent activity was defined as the index corresponding to the maximum derivative of the filtered ΔF/F_0_ trace of Crus I. The onset of plateau activity was determined as the first zero-crossing in the second derivative following the inflection point. Plateau duration was quantified by integrating over the distance or time for which activity remained above 20% ΔF/F_0_ during the pre-target interval.

### Mutual information and transfer entropy

We employed the Java Information Dynamics Toolkit to compute transfer entropy and mutual information using the Kraskov-Stögbauer-Grassberger estimator (Kraskov et al., 2004; Lizier, 2014). We used k = 4 nearest neighbors, normalized all traces, and used a maximum delay of 200 ms for transfer entropy. Mutual information and transfer entropy were calculated on a per-trial basis using the ΔF/F_0_ traces of Crus I and II in either 500 ms windows or a three-second interval preceding lick onset, respectively.

### Generalized linear model

We concatenated behavioral and task-related variables on a per-session basis and used the *glmfit* function in MATLAB to assess their contribution to the mean ΔF/F_0_ activity in Crus I and II (Engelhard et al., 2019). Predictor variables, whether continuous, event-based or whole-trial, were divided into four groups: body kinematics, reinforcement, pellet position and licks. Body kinematics included continuous predictors quantifying whole-body, eye, and whisker pad intensity rate, whole-body intensity acceleration, facial axis angle as well as snout and paw movement. Reinforcement included whole-trial variables such as binary trial outcome, hit rate in a ten-trial window, and a binary event variable marking a 50 ms window following the offset of a successful lick. Pellet position consisted of a single continuous variable representing the spatial distance of the stage from the midpoint of the mouse. Licks included a binary event variable marking frames in which the tongue was detected, as well as an event variable for the mean speed of each lick. Pellet position was normalized to a range of 0 to 1, peaking around the midpoint of the mice. All predictors, except binary variables, were z-scored. Random noise was added to binary and whole-trial variables to prevent ill-conditioning of matrices. The maximum variance inflation factor across all runs ranged from 2 to 3.7 (Median = 2.5), indicating low multicollinearity. We fit the models separately to the rest, pre-target and peri-target intervals using fifteen-fold cross-validation and trial weights to account for differences in trial length at different pellet speeds. Model fit was assessed by calculating R^2^ for each fold and then averaging across folds. Negative R^2^ values were set to zero. Relative contributions were calculated during cross-validation by systematically dropping each predictor group, refitting the reduced models, and subtracting the ratio of the R^2^ values of the reduced and full model from 1. Negative R^2^ values for the reduced models were set to 1 x 10^−6^ to prevent numerical instability. These contributions were then normalized across all predictor groups by dividing each by the total sum (Engelhard et al., 2019). The model was fit without cross-validation to obtain β-weights, and the absolute β-weights of individual predictors were summed for each predictor group. Finally, we multiplied the β-weights for each predictor group with its’ relative contribution. The resulting β coefficients reflect a predictor group’s unique contribution (in SDs) to ΔF/F_0_ activity independent of the contribution of all other predictors.

### Linear mixed-effects models

For trial-based analyses, we used the *lme* or *glme* functions in MATLAB to estimate the influence of fixed predictors on dependent variables while accounting for variability introduced by repeated measures from individual mice (7 levels) and, depending on the analysis, pellet speeds (5 levels). The mixed-effects structure, *R*^2^, *t*-statistics, degrees of freedom, and *p*-values are reported in the figure captions or main text. Categorical predictors included outcome (hit vs. miss), group (MI+ vs. MI−), day (early vs. late) and phase (baseline vs. CNO vs. saline). For binary dependent variables, we specified a binomial distribution and logit link function.

### Outlier removal

For the analysis of lick speed, any data points exceeding 3 SDs (221 mm/s) from the mean lick speed of all licks were excluded from individual analyses (∼ 0.1% of trials excluded).

### Data visualization

All figures were generated using GraphPad Prism version 10.3.1 for Windows (GraphPad Software).

### Statistical procedures

Test statistics are specified in the figure captions or the main text. All tests were two-tailed and statistical significance was defined at α = 0.05. Significance levels are indicated by * (p < 0.05), ** (p < 0.01), and *** (p < 0.001).

## Acknowledgements

We thank the OIST Animal Resources Section and Dr. Kazuo Mori for technical support, and Dr. Kiyoto Kurima for preparing plasmid DNA for AAV constructs. We are grateful to Profs. Kenji Doya, Sam Reiter and Kazumasa Tanaka as well as Dr. Soheil Keshmiri and members of the Optical Neuroimaging Unit at OIST for valuable comments on presentations of previous versions of this work. We thank the OIST Graduate University for generous financial support. This work was additionally supported by a JSPS Fellowship/KAKENHI grant (22J10169) awarded to M.E. and a KAKENHI grant (23H02454) awarded to B.K.

## Declaration of generative AI and AI-assisted technologies in scientific writing

During the preparation of this work, the authors used the large language model ChatGPT (OpenAI) to obtain suggestions for concise edits of parts of the abstract, introduction, and discussion. The authors reviewed and edited all content and take full responsibility for the content of the published article.

## Contributions

Conceptualization: C.G. and M.E.; Data collection: C.G.; Data analysis: C.G. and M.E.; Writing: C.G. and M.E.; Funding: M.E. and B.K. Supervision: B.K.

## Resource availability

### Lead contact

Requests for further information and resources should be directed to and will be fulfilled by any of the corresponding authors.

### Materials availability

This study did not generate new unique reagents.

### Data and code availability

Data and code are available from the lead contacts upon request.

## Author note

This research was funded by the Okinawa Institute of Science and Technology Graduate University (OIST) with subsidy funding from the Cabinet Office, Government of Japan. This research was additionally supported by JSPS/KAKENHI grant 22J10169 awarded to M.E. and KAKENHI grant 23H02454 awarded to B.K. The authors declare no conflicts of interest. Correspondence regarding this manuscript should be addressed to Cedric Galetzka, Mohamed Eltabbal, and Bernd Kuhn

---

**Figure S1:**
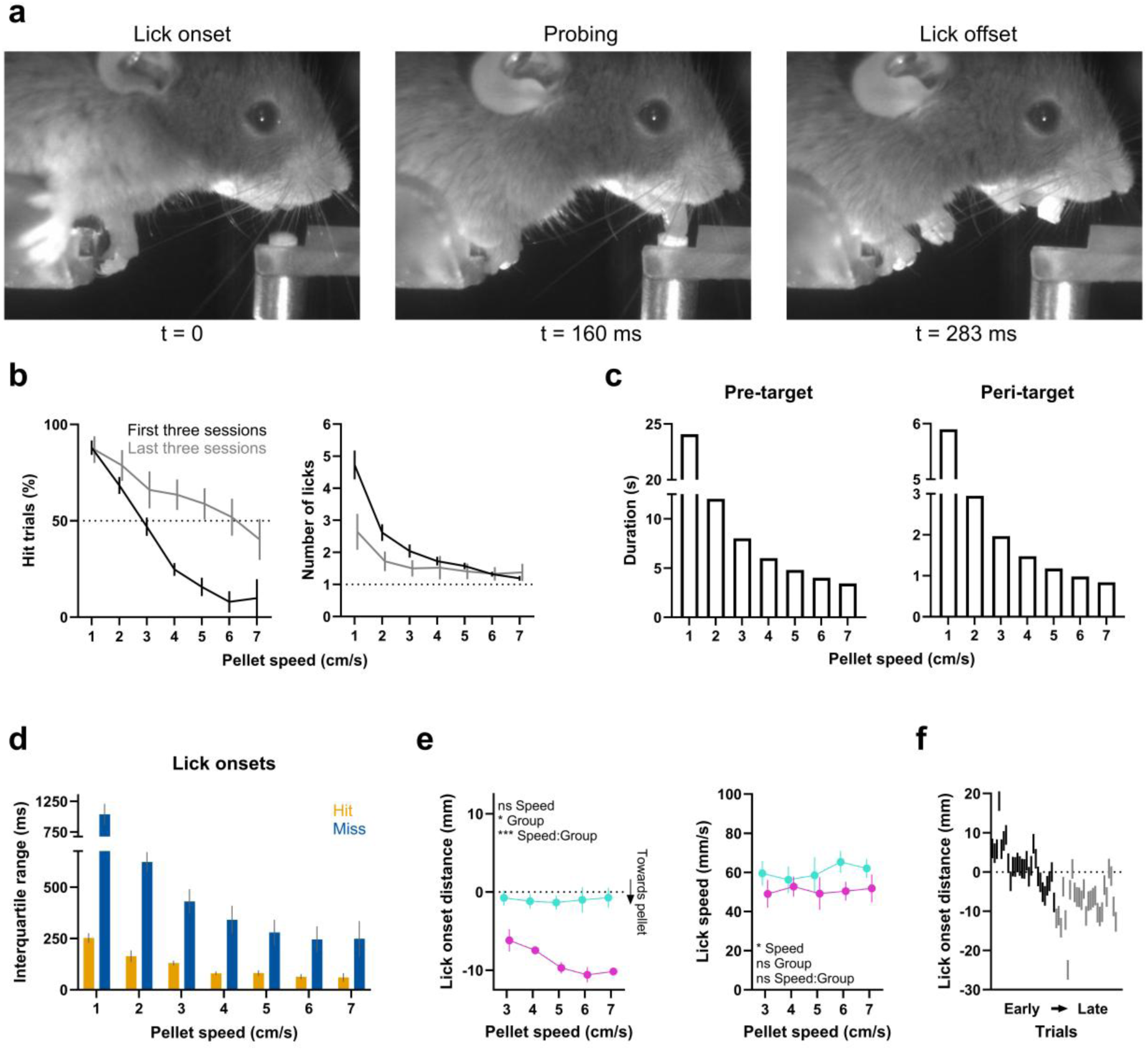
Performance, timing requirements, and successful lick parameters in MI+ and MI− mice. **(a)** Single video frames illustrating a representative lick during a hit trial, from lick onset to offset. **(b)** Left: Percentage of hit trials for each pellet speed during the first and last three sessions. Right: Mean number of licks. Data were computed for individual mice and then averaged across mice. Error bars denote the SEM (N = 7 mice). **(c)** Left: Bar plot of the duration of the pre-target interval at each pellet speed. Right: Duration of the peri-target interval. **(d)** Range of lick onsets for hit and miss trials at each pellet speed. Data shown were computed for individual mice and then averaged across mice. Error bars denote the SEM (N = 7 mice). **(e)** Left: Lick onset distance for pellet speeds 3-7 cm/s for MI+ and MI− mice during successful trials only. A linear mixed-effects model (*R*^2^ = 0.52) with pellet speed (3-7 cm/s) and group (MI+ vs. MI−) as fixed effects and mouse (7 levels) as random effect indicated a non-significant main effect of speed (t(409) = −0.326, p = 0.745), a significant main effect for group (*t*(409) = −2.293, *p* = 0.022), and a significant interaction between group and pellet speed (t(409) = −3.419, p < 0.001). Right: The same but for lick speed. A linear mixed-effects model (*R*^2^ = 0.35) revealed significant main effects of pellet speed (*t*(409) = 2.582, *p* = 0.01), no significant main effect for group (*t*(409) = −0.162, *p* = 0.871), and a marginally significant interaction between pellet speed and group (*t*(409) = −1.836, *p* = 0.067). Data was first averaged across early or late days for individual mice and then across mice. Error bars denote the SEM across mice (N = 4 MI+ and 3 MI− mice). **(f)** Lick raster of the first lick onsets in all trials of the same pellet speed from early to late learning for an example MI+ mouse.

**Figure S2:**
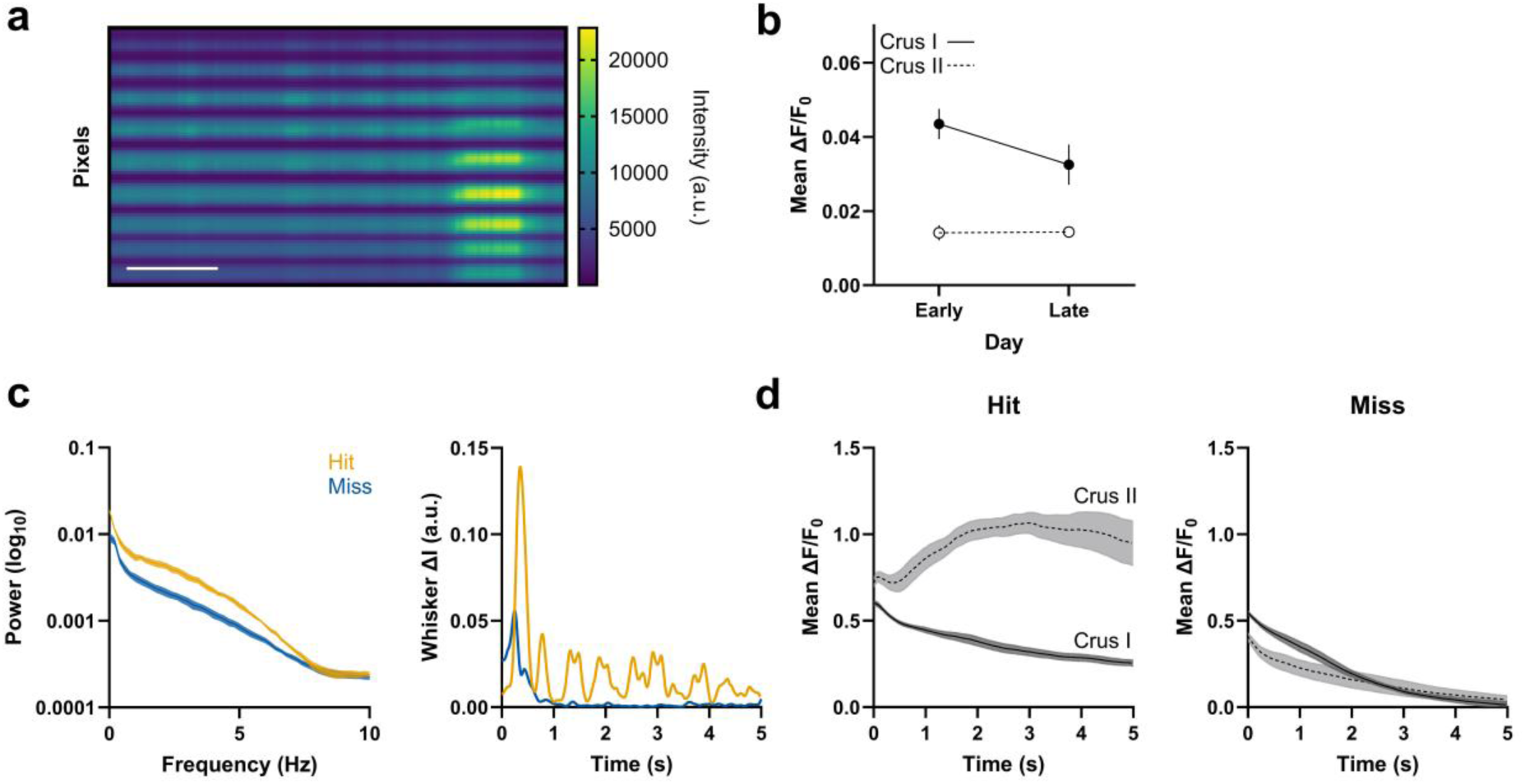
Interneuron activity in Crus II is elevated during orofacial movements associated with chewing. **(a)** Raw fluorescence intensity during the longest recorded interval at a pellet speed of 1 cm/s. Scale bar = 10 s; a.u. = arbitrary unit. **(b)** Mean ΔF/F0 during rest for Crus I and II. A linear mixed-effects model (*R*^2^ = 0.04) with day (early vs. late) and region (Crus I vs. Crus II) as fixed effects and mouse (7 levels) as random effect showed a non-significant main effect for day (*t*(2820) = −0.044, *p* = 0.965), a significant main effect for region (*t*(2820) = 5.199, *p* < 0.001) and a significant interaction between day and region (*t*(2820) = −2.219, *p* = 0.027). **(d)** Left: Power spectrum of the whisker pad intensity rate during the first five seconds of the post-target interval. Spectra were first computed for individual mice and then averaged across mice. Error bars denote the SEM over mice (N = 5 mice). Right: Whisker pad activity during the post-target interval for two representative hit and miss trials. ΔI = intensity rate. **(e)** Left: Mean ΔF/F_0_ activity during the first five seconds of the post-target interval for Crus I and II during hit trials. Right: Mean ΔF/F0 activity during miss trials. Traces were averaged across trials for individual mice and then across mice. Error bars denote the SEM over mice (N = 5 mice).

**Figure S3:**
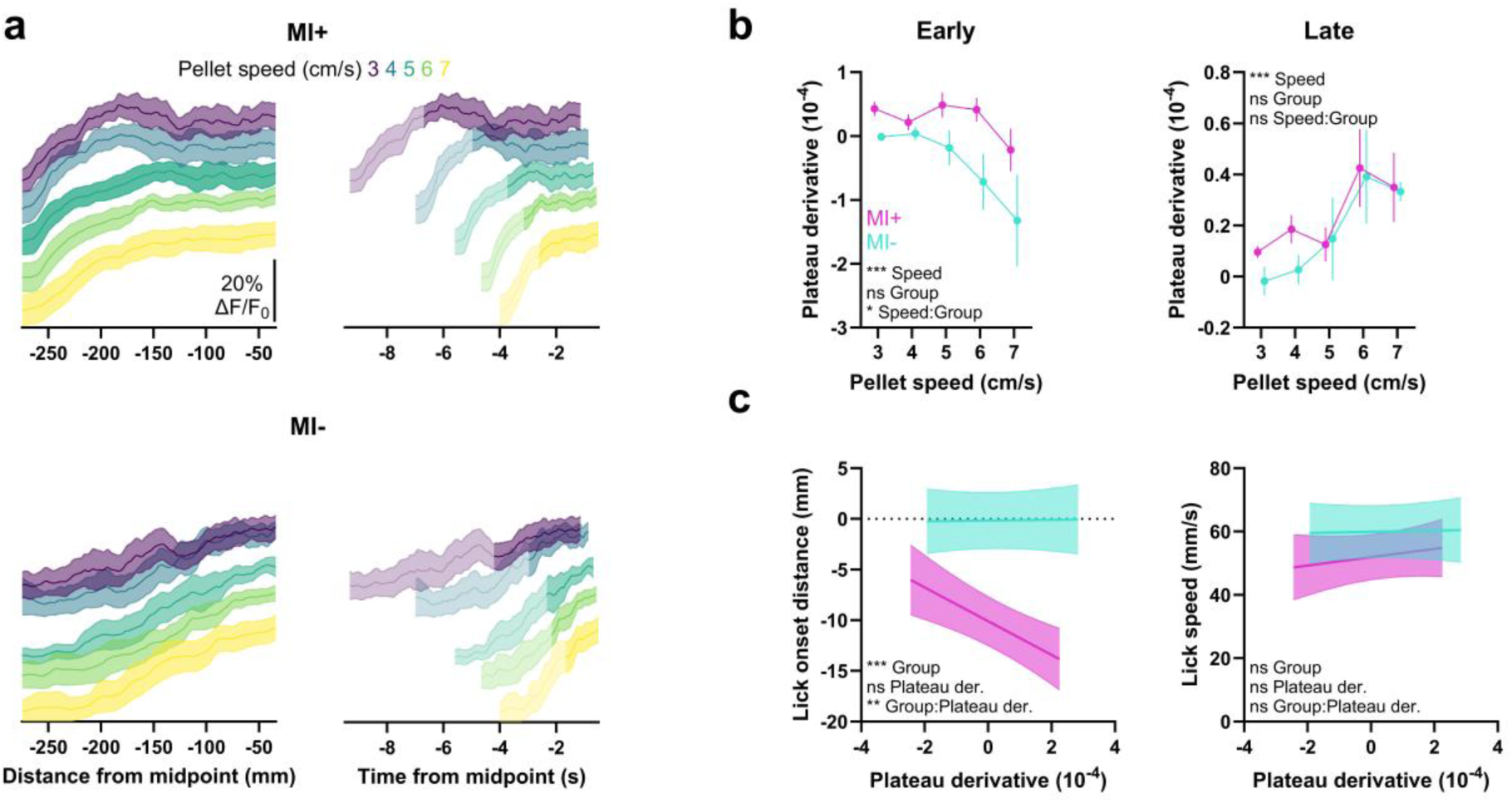
Persistent activity dynamics explain lick onset timing in MI+ mice. **(a)** Representative mean ΔF/F_0_ traces for an MI+ and MI− mouse during late learning. The same traces are plotted over distance or time from the midpoint. Shading denotes the 95% confidence interval. The opaque segments denote the portions of the traces included in the calculation of the plateau derivative, on average. Traces were vertically offset for visualization. **(b)** Left: Line plots of the mean plateau derivative as a function of pellet speed during early learning. A linear mixed-effects model (*R*^2^ = 0.09) with pellet speed (3-7 cm/s) and group (MI+ vs. MI−) as fixed effects and mouse (7 levels) as random effect indicated a significant main effect of pellet speed (*t*(498) = −3.477, *p* < 0.001), a non-significant main effect of group (*t*(498) = −0.394, *p* = 0.694) and a significant interaction between pellet speed and group (*t*(498) = 2.092, *p* = 0.037). Right: Mean plateau derivative during late learning. A linear mixed-effects model (*R*^2^ = 0.04) revealed a significant main effect of pellet speed (*t*(786) = 4.989, *p* < 0.001) and a non-significant main effect for group and interaction between pellet speed and group (all *p* > 0.05). Data shown was first averaged across trials. Error bars denote the SEM (N = 4 MI+ and 3 MI− mice). **(c)** Left: Lick onset distance as a function of the mean plateau derivative in MI+ and MI− mice during late learning. A linear mixed-effects model (*R*^2^ = 0.51) with group and plateau derivative as fixed effects and mouse as random effect showed a significant main effect of group (*t*(786) = −5.311, *p* < 0.001), a non-significant main effect for plateau derivative (*t*(786) = 0.098, *p* = 0.922), and a significant interaction between Group and plateau derivative (*t*(786) = −2.809, *p* = 0.005). Right: Lick speed in relation to the plateau derivative. A linear mixed-effects model (*R*^2^ = 0.2) revealed non-significant main effects of group and plateau derivative and a non-significant interaction between group and plateau derivative (all *p* > 0.05). Lines represent fitted regression lines from the linear mixed-effects models, with the shaded area indicating the 95% confidence intervals.

**Figure S4:**
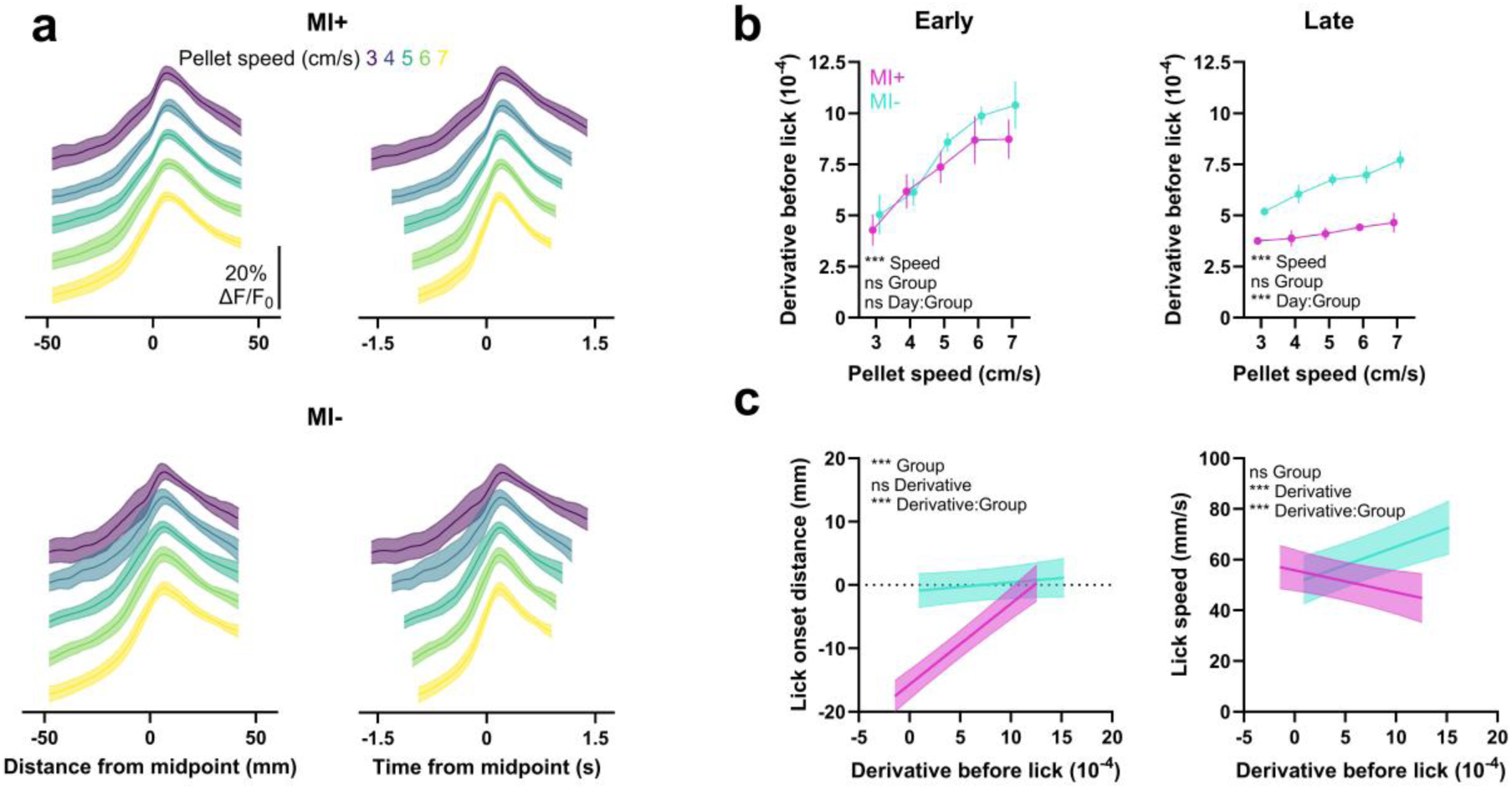
Sensory-related signals at the midpoint explain changes in lick speed in MI−mice. **(a)** Mean ΔF/F_0_ traces of Crus I for MI+ and MI− mice. The same traces are plotted over distance or time from the midpoint. Traces are vertically offset for visualization. Shading denotes the SEM (N = 4 MI+ and 3 MI− mice). **(b)** Left: Line plots of the mean derivative 250 ms before lick onset as a function of pellet speed during early learning. A linear mixed-effects model (*R*^2^ = 0.36) with pellet speed (3-7 cm/s) and group (MI+ vs. MI−) as fixed effects and mouse (7 levels) indicated a significant main effect of pellet speed (*t*(700) = 12.491, *p* < 0.001), a non-significant main effect of group and a non-significant interaction between pellet speed and group (all *p* > 0.05). Right: Mean derivative during late learning. A linear mixed-effects model (*R*^2^ = 0.31) revealed a significant main effect of pellet speed (*t*(839) = 8.933, *p* < 0.001), a non-significant main effect for group (*t*(839) = −0.905, *p* = 0.366), and a significant interaction between pellet speed and group (*t*(839) = −3.885, *p* < 0.001). Data shown was first averaged across trials. Error bars denote the SEM (N = 4 MI+ and 3 MI− mice). **(c)** Left: Lick onset distance as a function of the mean derivative before lick onset in MI+ and MI− mice during late learning. A linear mixed-effects model (*R*^2^ = 0.58) with group and derivative as fixed effects and mouse as random effect showed a significant main effect of group (*t*(839) = −7.968, *p* < 0.001), a non-significant main effect for derivative (*t*(839) = 1.284, *p* = 0.2) and a significant interaction between group and derivative (*t*(839) = 7.134, *p* < 0.001). Right: Lick speed in relation to the derivative. A linear mixed-effects model (*R*^2^ = 0.24) revealed a non-significant main effect of group (*t*(839) = 0.8, *p* = 0.424), a significant main effect of derivative (*t*(839) = 4.167, *p* < 0.001) and a significant interaction between group and derivative (*t*(839) = −4.6629, *p* < 0.001). Lines represent fitted regression lines from the linear mixed-effects models, with the shaded area indicating the 95% confidence interval.

